# Heat stress induces ferroptosis in a photosynthetic prokaryote

**DOI:** 10.1101/828293

**Authors:** Anabella Aguilera, Federico Berdun, Carlos Bartoli, Charlotte Steelheart, Matías Alegre, Graciela Salerno, Gabriela Pagnussat, María Victoria Martin

## Abstract

Ferroptosis is an oxidative iron-dependent form of cell death recently described in eukaryotic organisms like animals, plants and parasites. Here we report that a similar process takes place in the cyanobacterium *Synechocystis* sp. PCC 6803 in response to heat stress. After a heat shock, *Synechocystis* cells undergo a cell death pathway that can be suppressed by canonical ferroptosis inhibitors or by external addition of calcium, glutathione or ascorbic acid. Moreover, as described for eukaryotic cells ferroptosis, this pathway is characterized by an early depletion of antioxidants, and by lipid peroxidation. As in general prokaryotes membranes contain poorly oxidizable saturated or monounsaturated lipid molecules, it was thought that they were not susceptible to ferroptosis. Interestingly, cyanobacteria contain thylakoid membranes that are enriched in polyunsaturated-fatty-acid-containing phospholipids, which might explain their sensitivity to ferroptosis. These results indicate that all of the hallmarks described for eukaryotic ferroptosis are conserved in photosynthetic prokaryotes and suggest that ferroptosis might be an ancient cell death program.

**Summary:** Aguilera et al, show that ferroptosis, an oxidative and iron-dependent form of regulated cell death, plays an important role in the cyanobacterium *Synechocystis* sp. PCC 6803 in response to heat stress.

## Introduction

In contrast with Accidental Cell Death (ACD), Regulated Cell Death (RCD) relies on a tightly modulated molecular machinery that involves signalling cascades and defined effectors (Galluzzi et al., 2018). In eukaryotes, RCD plays a critical role in essential physiological programs such as embryonic development, differentiation, fertilization, tissue renewal and immune responses, being generally referred to as Programmed Cell Death (PCD) (Hakem et al., 1998; Lindsten et al., 2000; Van Hautegem et al., 2015; Yoshida et al., 1998). RCD can also occur when responses to perturbations of the intracellular or extracellular microenvironment fail, as an ultimate attempt to maintain homeostasis (Galluzzi et al., 2016).

Different types of RCD have been described in eukaryotes (Melino et al., 2005; Minina et al., 2013). Among them, ferroptosis was recently reported as an oxidative, iron-dependent form of RCD associated with lipid peroxidation and ROS accumulation present in animals, plants and protozoan parasites (Bogacz and Krauth-Siegel, 2018; Conrad et al., 2018; Dangol et al., 2018; Distéfano et al., 2017; Dixon et al., 2012; Stockwell et al., 2017).

Although less studied and understood, RCD also occur in eukaryotic microorganisms and at least in some prokaryotes (Allocati et al., 2015; Bayles, 2014). Cyanobacteria are widely distributed Gram-negative bacteria that are capable of plant-like oxygenic photosynthesis (Whitton, 2012). In particular, cyanobacteria are important components of phytoplankton communities, contributing to a substantial fraction of the global primary production and are a crucial source of atmospheric oxygen (Franklin, 2014). Despite its ecological and biogeochemical significance, the nature of RCD in cyanobacteria is still not completely understood (Bidle, 2016). Evidence has accumulated on controlled cell-death mechanisms triggered under unfavorable environmental conditions, such as nutrient deprivation, high-light-associated oxidative stress or osmotic stress that are indistinctly termed as PCD, apoptotic-like or necrotic-like death. Such cell-death pathways involve morphological changes, accumulation of reactive oxygen species (ROS), DNA laddering, loss of plasma membrane integrity, and the coordinated participation of redox enzymes, metabolites and caspase-like proteases (Bidle, 2015; Durand et al., 2016; Spungin et al., 2018; Swapnil et al., 2017; Zhou et al., 2018). RCD has been proposed to optimize differentiation, dynamics and colony fitness in cyanobacteria (Bar-Zeev et al., 2013; Meeks and Elhai, 2002). In addition, microscopic analyses showed that cyanobacteria symbiotically associated with the water-fern *Azolla microphylla* can follow different RCD pathways with characteristics of metazoan apoptosis, autophagy, necrosis and autolysis (Zheng et al., 2013).

In this work, we examined whether ferroptosis could be relevant to the model cyanobacterium *Synechocystis* sp. PCC 6803 (hereafter *Synechocystis*) under abiotic stresses. We found that in response to heat stress (50°C), *Synechocystis* follows a cell death program that shows biochemical and morphological features that resembles eukaryotic ferroptosis. This cell death is dependent on iron availability, lipid peroxidation, and is inhibited by canonical ferroptosis inhibitors. Moreover, cell death is characterized by depletion of glutathione (GSH) and ascorbic acid (AsA) and can be prevented by GSH or AsA addition. These results not only suggest that ferroptosis might be an ancient cell death program. As chloroplasts originated from the endosymbiosis of cyanobacterial-like organisms, the presence of this iron-dependent oxidative cell death pathway in cyanobacteria also points to the evolutionary origin of the chloroplasts role during plant ferroptosis (Distéfano et al., 2017).

## Results and discussion

### Ferroptosis inhibitors prevent the regulated cell death pathway triggered by a 50°C-heat shock in cyanobacteria

Heat stress is known to cause cell death in *Synechocystis* (Suginaka et al., 1999). That pathway, triggered by a 50°C heat shock (HS), is characterized by a strong decrease in GSH levels, a feature commonly observed in eukaryotic ferroptosis (Conrad et al., 2018; Distefano et al., 2017; Dixon et al., 2012). To examine whether ferroptosis plays a role during stress-induced cell death in the cyanobacterium, cells were exposed to H_2_O_2_ or to high temperatures at 50°C, or at 77°C (to trigger unregulated necrosis), in the presence of two canonical ferroptosis inhibitors: the lipophilic antioxidant ferrostatin-1 (Fer-1) or the membrane-permeable iron chelator ciclopirox olamine (CPX). Cell death was assessed by three different methodologies following NCCD recommendations (Kroemer et al., 2009): drop tests to measure cell survival (Fig. 1a), fluorescence microscopy using SYTOX® Green as a cell death marker (Fig 1b and S1a) and FDA fluorescence quantification by flow cytometry (Fig. 1c). All experiments indicated that cell death triggered by 50°C is significantly prevented by Fer-1 and by CPX (Fig. 1, a-c), suggesting that the 50°C-HS triggers a ferroptotic cell death pathway. In contrast, neither Fer-1 nor CPX were able to prevent cell death triggered by a higher temperature (77°C, aimed to promote an unregulated necrosis) or by H_2_O_2_, already known to promote cell death in a regulated way in the cyanobacterium *Microcystis aeruginosa* (Ding et al., 2012)(Fig. 1 c and S1b). These results indicate that the response after the 50°C-HS is specific.

**Figure 1.**
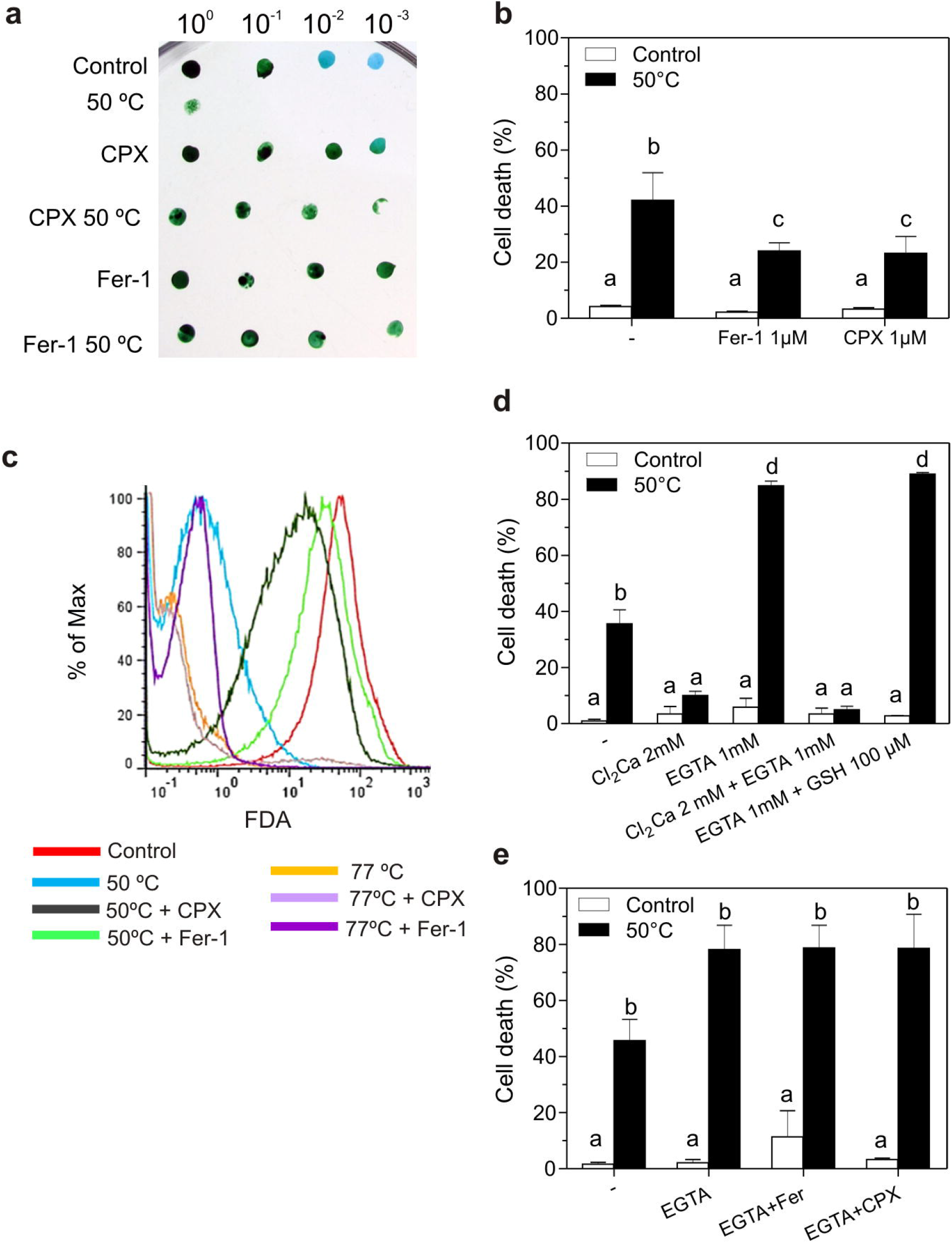
Ferroptosis inhibitors and calcium prevent cell death induced by a 50°C heat shock in *Synechocystis* sp. PCC 6803. **(a)** Viability of *Synechocystis* sp. PCC 6803 cells exposed to 50°C for 10 min preincubated with DMSO (−), with Fer-1 (1 μM) or with CPX (1 μM) for 24 h, was tested via a drop test on BG11 agar plates. The effect of each treatment was verified by three independent experiments. **(b)** HS-induced cell death in *Synechocystis* sp. PCC 6803 preincubated with DMSO, Fer-1 or CPX for 24h. **(c)** Cell viability assessed by flow cytometry using the FDA probe after inducing cell death after a treatment of 50°C or 77°C for 10 min in cultures preincubated with DMSO, Fer-1 or CPX for 24h. **(d)** Cell death assessed in *Synechocystis* sp. PCC 6803 preincubated with Cl_2_Ca, with EGTA, or with Cl_2_Ca-EGTA before inducing cell death by treating cells at 50°C for 4 h. (e) Cell death assessed in *Synechocystis* sp. PCC 6803 cells preincubated with EGTA and then with Fer-1 or CPX before inducing cell death by treating cells at 50°C for 4 h. (b, d, e) Cell suspensions were stained with SYTOX Green, examined and counted under light and fluorescence microscopy. SYTOX-positive cells were interpreted as dead cells. Box plots are from at least three independent experiments. Plots with different letters denote statistical difference (two-way ANOVA in GLM, P < 0.05).

The role of calcium in ferroptosis is still a matter of debate. While the calcium chelator ethylene glycol-bis (b-aminoethylether)-N,N,N’,N’-tetraacetic acid (EGTA) was found to block ferroptosis in plants (Distéfano et al., 2017), chelation of extracellular calcium does not prevent cell death in response to glutathione depletion in human cancer cells (Dixon et al., 2012). Another report, however, indicate that extracellular calcium influx is required for cell death downstream of glutathione depletion in mammalian neuronal-like HT22 cells (Henke et al., 2013). These different behaviours suggests that calcium requirement could be cell-type specific. To test the effect of calcium, we exposed *Synechocystis* cells to a HS at 50°C in the presence of CaCl_2_, EGTA or both. After a 4 h treatment, about 50% of 50°C-treated cells were dead. Surprisingly, when co-treated with CaCl_2_, only ∼10% of the 50°C-treated cells died, a value comparable to the control (cells that were not exposed to HS) (Fig. 1d). On the other hand, ∼80% of the cells were found dead when treated with EGTA and less than ∼10% were dead when EGTA and CaCl_2_ were added together. These results suggest that extracellular calcium can prevent cell death induced by 50°C-HS (Fig. 1 d). However, neither Fer-1 nor CPX were able to prevent cell death in the presence of EGTA (Fig. 1 e). These findings suggest that calcium depletion could be involved in HS-induced cell death as a very early event, and are in agreement with previous results showing that exogenous Ca^++^ supplementation improve the tolerance of *Anabaena* sp. PCC 7120 to HS (Tiwari et al., 2016).

AsA and GSH are two key components of the antioxidant machinery of eukaryotic cells (Foyer and Noctor, 2011; Aquilano et al., 2014). In mammalian cells, inactivation of the selenoenzyme glutathione peroxidase (GPX4) through GSH depletion results in an overwhelming lipid peroxidation that triggers ferroptosis (Dixon et al., 2012; Skouta et al., 2014). Similarly, ferroptosis in plants is characterized by the early depletion of GSH and AsA (Distéfano et al., 2017). In addition, GSH depletion is required for erastin-induced hypersensitive response in rice-*Magnaporthe oryzae* interaction (Dangol et al., 2018). *Synechocystis* presents both compounds in different proportions (Suginaka et al., 1999). As described for eukaryotic systems, the exposure of *Synechocystis* cells to 50°C results in a decline of GSH and AsA total contents (Fig. 2, a and b). These effects were not prevented by the addition of Fer-1 or CPX, suggesting that GSH and AsA depletion might be an early event, as seen in animal and plant cells (Dixon et al., 2012; Distéfano et al., 2017). The redox status of glutathione (% of oxidized GSSG), did not show a clear association with the treatments and did not show differences with the addition of Fer and CPX under normal temperature conditions. On the other hand, oxidized AsA content was very low or almost undetectable. Notably, cell death was prevented by pre-incubation with GSH and AsA, suggesting that GSH and AsA depletion is required for HS-induced cell death in *Synechocystis* (Fig. 2 c).

**Figure 2.**
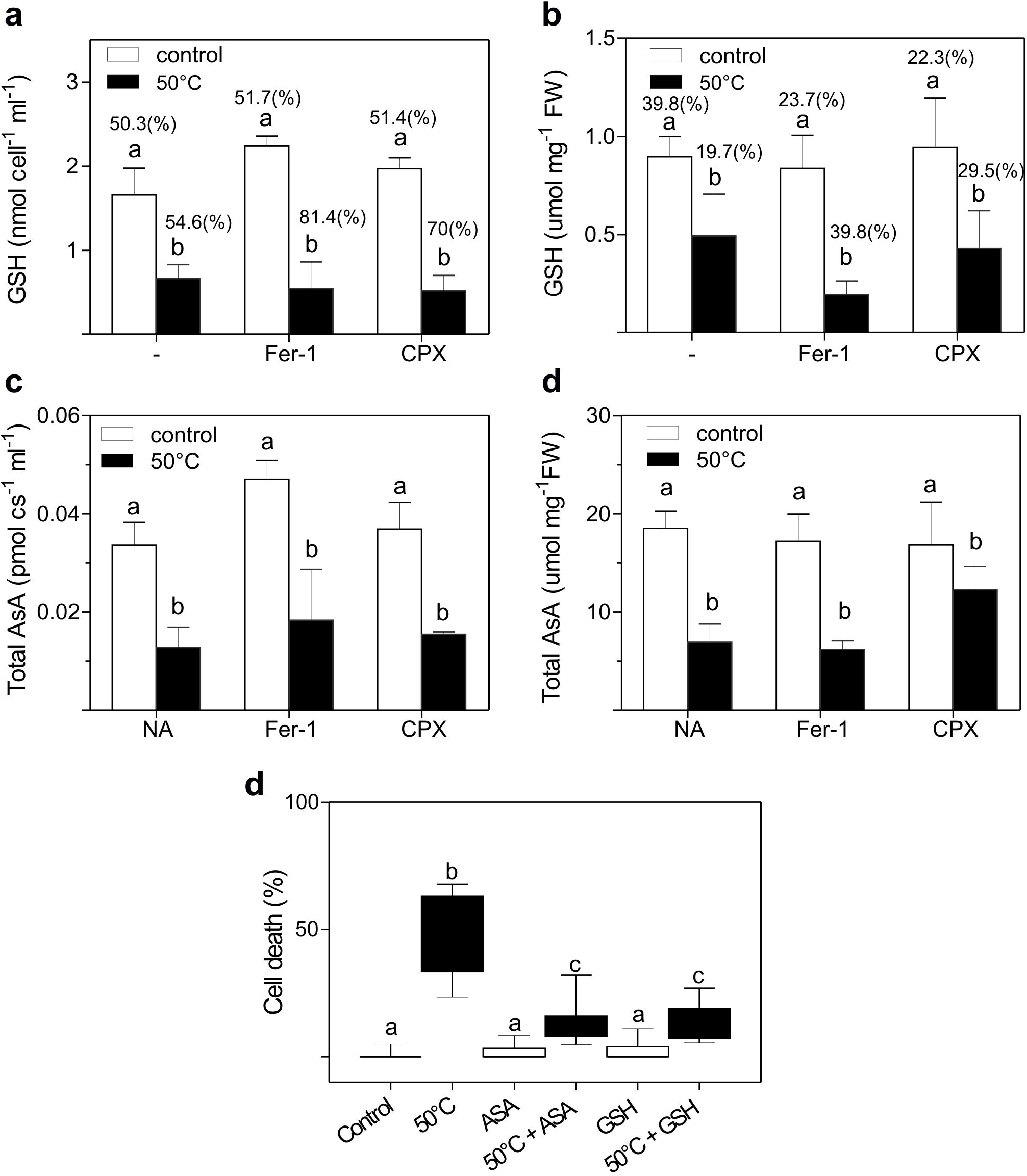
Cell death triggered by 50°C induces GSH and AsA depletion in *Synechocystis* sp. PCC 6803. GSH levels **(a)** per living cell. **(b)** per mg of fresh weight. Total AsA content **(c)** per living cell **(d)** per mg of fresh weight, measured in *Synechocystis* sp. PCC 6803 after treating cells at 50°C for 4 h. In each case, preincubation with DMSO (−), Fer-1 (1 μM) or CPX (1 μM) 24 h before HS is indicated. **(e)** Cell death induced by a 50°C treatment is prevented by GSH (100 μM) or AsA (1 μM) addition 24 h before HS. Cell suspensions were stained with SYTOX Green, examined and counted by light and fluorescence microscopy. Sytox-positive cells were interpreted as dead cells. (a and b) Percentage indicate redox status (% of oxidized). Different letters denote statistical difference (two-way ANOVA, P < 0.05). (d) Box plots with different letters denote statistical difference (two-way ANOVA in GLM, P < 0.05). Data are from three independent experiments.

### 50°C heat shock-induced cell death involves rapid accumulation of cytosolic and lipid ROS in *Synechocystis*

In eukaryotic organisms, cytoplasmic ROS and lipid ROS accumulation play a central and essential role in the ferroptotic pathway. The oxidative burst observed in eukaryotic cells undergoing ferroptosis is dependent on iron as it is prevented by iron chelators (Bogacz and Krauth-Siegel, 2018; Distéfano et al., 2017; Dixon et al., 2012). In order to investigate if a similar process occurs in *Synechocystis*, cytosolic ROS and lipid ROS accumulation were measured after inducing cell death by 50°C-HS. The accumulation of cytosolic ROS was measured by using the probe H_2_DCFDA, while the fluorescence intensity of the lipid peroxidation probe C11-BODIPY was monitored as an indicator of lipid ROS content. A maximum of cytosolic ROS was detected 1 h after 50°C-HS cell death induction (Fig. 3a). This accumulation of cytosolic ROS was prevented by incubation with Fer-1 or CPX (Fig. 3a). The accumulation of lipid peroxides reached a maximum 3 h after the 50°C-HS and was suppressed by ferroptosis inhibitors (Fig. 3b).

**Figure 3.**
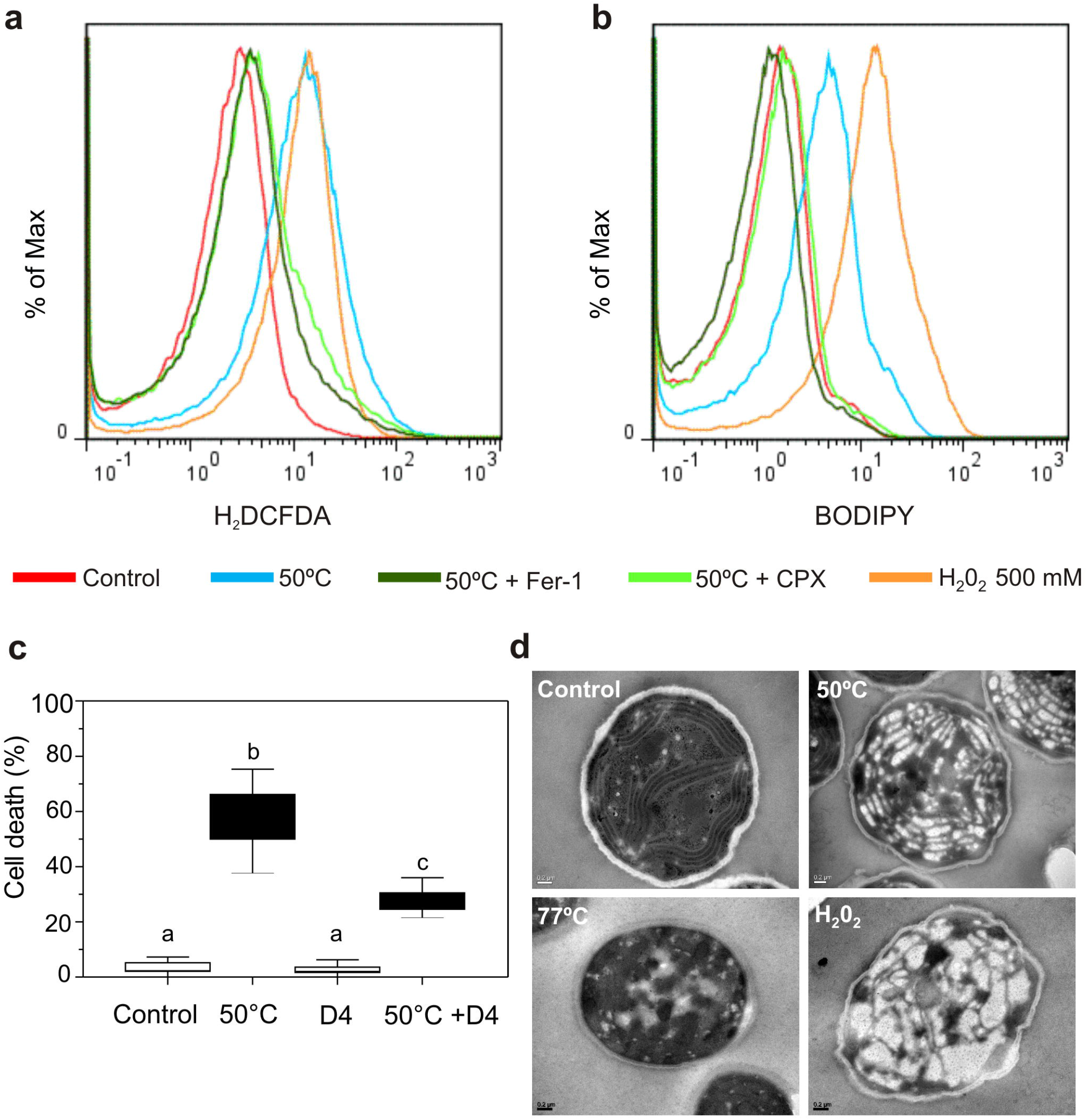
50°C treatment triggers morphological changes and accumulation of ROS and lipid ROS that are prevented by ferroptosis canonical inhibitors. **(a, b)** Cytosolic and lipid ROS levels were assessed by flow cytometry at 1 h and 3 h, respectively, after a 10 min 50°C treatment using H_2_DCFDA and BODIPY. Cultures were preincubated with DMSO, Fer-1 (1 μM) or CPX (1 μM) for 24 h as indicated. Aliquots treated with 500 mM H_2_O_2_ were used as positive controls for ROS content and oxidized lipids production. **(c)** Cell death after 50°C is prevented by D-PUFAs (D4). Cultures were preincubated with DMSO or with 50 μM D4-linoleate for 24 h. Cell death was induced by treating cells at 50°C for 4 h. **(d)** TEM micrographs of *Synechocystis* sp. PCC 6803 cells. Cultures were treated with DMSO (control), 50°C, or 77°C for 10 min or with H_2_O_2_ 10 mM for 1 h. **(c)** Cell suspensions were stained with SYTOX Green, examined and counted by light and fluorescence microscopy. SYTOX-positive cells were interpreted as dead cells. Box plots are from three independent experiments. Plots with different letters denote statistical difference (two-way ANOVA in GLM, P < 0.05).

In human and plant cells, lipid peroxidation during ferroptosis was shown to involve peroxidation of polyunsaturated fatty acids (PUFAs) at the bis-allylic position. Pre-treatment with PUFAs containing the heavy hydrogen isotope deuterium at the site of peroxidation (D-PUFAs) is able to prevent ferroptosis, both in human cancer cells and in plant root cells (Yang et al., 2016; Distéfano et al., 2017). Likewise, we observed a clear protective effect of D-linoleate (D-Lin, 16 h) against 50°C-HS in *Synechocystis* sp. PCC 6806 (Fig. 3c). Altogether, these results indicate that cyanobacterial cells exposed to a 50°C-undergo an oxidative, iron-dependent form of cell death remarkably similar to ferroptosis in eukaryotic cells (Dixon et al., 2012; Distéfano et al., 2017; Bogacz and Krauth-Siegel, 2018; Dangol et al., 2018).

### *Synechocystis* shows thylakoid membranes alteration after heat shock

To gain insight into the morphological changes taking place in our cell death model, *Synechocystis* sp. PCC6803 cells treated with H_2_O_2_ or subjected to 50°C or to 77°C were studied using transmission electron microscopy (TEM) (Fig.3d). The distinctive morphological features of 50°C treated cells involved the loss of the thylakoids membrane integrity, with evident non-electron dense zones, and vesiculation (Fig. 3d). Notably, disruption of the thylakoid membranes and vesiculation are already known to occur after a heat treatment in plants (Ristic et al., 2007) and in the unicellular eukaryotic algae *Chlorella saccharophila* during cell death induced by heat (Zuppini et al., 2007). Remarkably, thylakoid membranes are enriched in polyunsaturated-fatty-acid-containing phospholipids (PUFA-PLs), which are the lipids most susceptible to peroxidation during ferroptosis (Sharathchandra and Rajashekhar, 2011; Strahl and Errington, 2017). On the other hand, cells exposed to 77°C showed a reduction of the cellular volume and electron dense regions, while the thylakoid membranes were not distinguishable (Fig. 3d). Cells treated with H_2_O_2_ showed cytoplasm vacuolation (Fig. 3d), a characteristic previously observed in *Microcystis aeruginosa* during cell death induced by high concentrations of H_2_O_2_ (Ding et al., 2012)

### No caspase-like activity is induced after 50°C-HS in *Synechocystis*

Metacaspases are cysteine specific proteases evolutionarily related to metazoan caspase proteases (Spungin et al., 2018) implicated in stress response and cell death in bacteria and phytoplankton. Caspase-like activities have been reported in *Microcystis aeruginosa* after treatments with pyrogallic acid, high light intensity or low phosphorus availability (He et al., 2016; Lu et al., 2017). *Trichodesmium* spp. shows an increase in caspase-like activities under low iron and high-light stress (Asplund-Samuelsson et al., 2012; Berman-Frank et al., 2004, 2007; Spungin et al., 2018).

Only one gene encoding a metacaspase from the cyanobacterial Family β (cbMC β) (Jiang et al., 2010) is present in the *Synechocystis* genome. Although it has been suggested that cbMC β might be catalytic inactive (Klemencic and Funk, 2018), the gene is highly expressed (Klemencic et al., 2019). To investigate whether caspase activity increases after cell death induction, caspase activity was directly assessed by using the CellEvent caspase-3/7 green detection reagent in *Synechocystis* cells exposed to HS or to an oxidative stress. Caspase-3/7 signals did not increased after 50°C, 77°C or after exposing the cells to H_2_O_2_ (Fig. S1 e) indicating that a caspase-like activity is not involved in the cell death processes induced by these triggers, in accordance with previous reports in animal cells (Dixon et al., 2012).

### Molecular mechanisms governing ferroptosis in *Synechocystis*

To gain insight into the regulation of several candidate genes that might be involved in cyanobacteria ferroptosis, we examined their expression after exposing *Synechocystis* cells to a 50°C-HS, pre-incubated or not with Fer-1 or CPX (Table 1). The genes analyzed include:

-genes involved in cyanobacterial GSH synthesis *(gshA* an *gshB)* and GSH catabolism (*ggt*) (Narainsamy et al., 2016); *gpx1* and *gpx2* orthologues to human GPX4, which is essential for ferroptosis in animals (Friedmann Angeli et al., 2014; Imai et al., 2017). Three genes, *gshA, ggt* and *gpx2* were up-regulated after 50°C exposure and only *gpx2* was down-regulated. On the other hand, *gpx1* and *gshB* were found to be poorly expressed in the analyzed conditions. Only *gpx2* showed a clear response to HS, an induction of about 10-fold that was prevented by CPX (Table 1).
-genes involved in Fe metabolism such as ferric and ferrous iron transporters (*futA, futB, futC* and *feoB*) and the *isiA* gene, that encodes a chlorophyll-binding protein inducible by high light (Havaux et al., 2005). *feoB* was highly up-regulated after the 50°C-HS, which was enhanced by pre-treatment with Fer-1 (Table 1).
-genes related to the heat shock regulon involved in heat stress (*groES, groEL*) (Rajaram et al., 2014). As can be observed in Table S1, *groES* and *groEL* were up-regulated after HS and that was not prevented by pre-treatment with ferroptosis inhibitors (Table 1).

**Table 1.** Changes in expression of candidate genes that might be associated with ferroptosis in *Synechococcus* sp. PCC 6803 30 min after a 50°C treatment for 1 h. The levels of mRNA are expressed as a fold-change ratio between the different conditions and control.

| Gene | 50°C | 50°C +Fer-1 | 50°C + CPX |
| --- | --- | --- | --- |
| *groEL1* | 18.9 ± 9.5 | 17.9 ± 9.1 | 12.1 ± 2.5 |
| *groES* | 13.2 ± 7.5 | 13.3 ± 1.1 | 13.9 ± 1.2 |
| *gshA* | 11.7 ± 0.6 | 34.0 ± 7.0 | 11.3 ± 5.1 |
| *gshB* | 0.9 ± 0.0 | 2.3 ± 0.7 | 0.4 ± 0.4 |
| *ggt* | 4.00 ± 0.9 | 8.5 ± 4.6 | 3.0 ± 1.5 |
| *gpx1* | 0.3 ± 0.2 | 0.2 ± 0.1 | 0.2 ± 0.1 |
| *gpx2* | 10.0 ± 0 | 10.0 ± 0 | n.d. |
| *feoB* | 42.3 ± 12.1 | 79.7 ± 17.0 | 25.0 ± 9.5 |
| *furA* | 0.6 ± 0.1 | 1.0 ± 0.1 | 0.5 ± 0.3 |
| *futA* | 4.3 ± 0.6 | 10.7 ± 5.7 | 3.0 ± 1.0 |
| *futB* | 2.8 ± 0.7 | 7.4 ± 3.4 | 2.2 ± 0.9 |
| *futC* | 0.7 ± 0.0 | 2.2 ± 1.5 | 0.5 ± 0.4 |

The analysis of the *Synechocystis* genome and data reported in previous studies, allowed us to identify several components of the eukaryotic ferroptosis pathway that are present in cyanobacteria:

Selenium-containing glutathione peroxidase 4 (GPX4) is a key regulator of eukaryotic ferroptosis (Yang et al., 2014; Seibt et al., 2019). The genome of *Synechocystis* encodes two GPX-like proteins annotated as GPX1 (slr1171) and GPX2 (slr1992), with high similarity to higher plants and mammals GPXs that do not contain selenium (Gaber et al., 2001). *In vitro* experiments have shown that recombinant GPX-like proteins of *Synechocystis* can use unsaturated fatty acid hydroperoxides or alkyl hydroperoxides as an acceptor, but they are unable to utilize GSH as an electron donor (Gaber et al., 2001). Cyanobacteria have multiple detoxifying enzymes to manage oxidative stress, such as glutathione peroxidases and peroxiredoxins (Johnson and Hug, 2019) that might also be involved in this response.

As noted above, eukaryotic ferroptosis is highly linked to GSH metabolism and content (Dixon and Stockwell, 2019). In cyanobacteria, GSH is a key antioxidant involved in the protection against several ROS species (Latifi et al., 2009). In this work, we showed that cell death induced by 50°Cwas preceded by the depletion of the antioxidants GSH and AsA (Fig. 2, a and b), as seen in mammalian cells and root hairs of *A. thaliana* (Yang and Stockwell, 2016; Distéfano et al., 2017; Dixon and Stockwell, 2019). Moreover, addition of GSH is able to prevent the cell death triggered by HS (Fig. 2 c), as reported in mammalian cells and *A. thaliana* (Dixon et al., 2012; Distéfano et al., 2017).

Long-chain acyl-CoA synthetase-4 (ACSL4) modulate the metabolic fates of PUFAs in human and mouse cells and deletion of the *ACSL4* gene causes resistance to ferroptosis in mammals (Conrad et al., 2018). The *Synechocystis* genome contain s sequence that encodes a putative acyl-acyl carrier protein synthetase (SynAas, Slr1609) that is a homologue of the *A. thaliana* long chain acyl-CoA synthetase 9 (LACS9). SynAas specifically uses acyl-acyl carrier protein as a co-substrate to recycle free fatty acids and it is also involved in the transfer of free fatty acids across membranes by vectorial acylation (Kaczmarzyk and Fulda, 2010; von Berlepsch et al., 2012). The oxidation of phospholipids containing PUFAs in one of the three essential hallmarks that define ferroptosis (Dixon and Stockwell, 2019). PUFAs are generally present in cyanobacterial membranes and thylakoid membranes are specifically enriched in PUFA-PLs. Several stress conditions such as high temperature, high light or algaecide exposure have been shown to trigger the generation of lipid peroxides (Maeda et al., 2005; Latifi et al., 2009; Lee et al., 2018, Allakhverdievet al. 1999; Singh et al., 2002). As in mammalian cells and root hairs of *A. thaliana* (Yang et al., 2016; Distéfano et al., 2017), the death in *Synechocystis* cells exposed to heat shock was prevented by supplementation with D-PUFAs (Fig. 3 c). PUFAs oxidation by lipoxygenases (LOXs; encoded by the ALOX genes) is required for ferroptotic cell death in mammals cells (Yang et al. 2016). Some lipoxygenase sequences have been detected in cyanobacteria such us from the genus *Anabaena, Nostoc, Microcystis*, and *Synechococcus*, and a mini-lipoxygenase has been biochemically characterized in *Cyanothece* (Andreou et al., 2010; Hansen et al., 2013). However, the biological role of bacterial lipoxygenases remains to be elucidated (Hansen et al., 2013).

Vitamin E (alpha-tocopherol) has been reported to prevent ferroptosis in human and mouse cells (Conrad et al., 2018; Hinman et al., 2018). Alpha-tocopherol is synthesized by some cyanobacteria and remarkably, it has a role protecting *Synechocystis* from lipid peroxidation and high light stress (Maeda et al., 2005).

Ferroptosis has also been described as an autophagy-dependent process (Zhou et al., 2019). While autophagy is present in almost all eukaryotes, little is known about whether this process occurs in cyanobacteria. Most prokaryotes kept three or four homologs of autophagy genes, but cyanobacteria carry approximately ten homologs. Notably, it was reported that endosymbiotic cyanobacteria present in the fern *Azolla microphylla* exhibit autophagy-like cell death that is characterized by a gradual condensation and degradation of the cytoplasm and high levels of vacuolization (Zheng et al., 2013). Accordingly, high levels of vacuolization were observed in *Synechocystis* cells after exposure to 50°C for 10 min (Fig. 3 d). These observations suggest that autophagy might also be part of the ferroptotic mechanism in cyanobacteria although more studies are needed to establish if that is the case.

Based on the results obtained in this study, we summarized our current understanding of the pathways leading to ferroptosis in the prokaryote *Synechocystis* (Fig. 4). Concisely, the pathway involves early GSH depletion following a HS, which leads to lipid peroxidation, mostly of polyunsaturated fatty acids from thylakoid and plasma membranes. Canonical ferroptosis inhibitors like the iron chelator ciclopirox CPX or the lipophilic antioxidant Fer-1, prevent the accumulation of toxic lipid and cytosolic ROS, inhibiting ferroptotic cell death. Mechanistically, this pathway shows remarkable similarities to plant and mammalian ferroptosis. Interestingly, thylakoid membranes seem to be involved in this process, possibly providing PUFAs that undergo peroxidation. Thylakoid membranes are enriched in PUFAs, which differentiates cyanobacteria from other prokaryotes, which in general carry membranes containing saturated or monounsaturated lipids (Sharathchandra and Rajashekhar, 2011). This fact can also be related to previous results obtained in the model plant *A. thaliana*, where active chloroplasts were shown to contribute to ferroptotic cell death in leaves (Distéfano et al., 2017).

**Figure 4.**
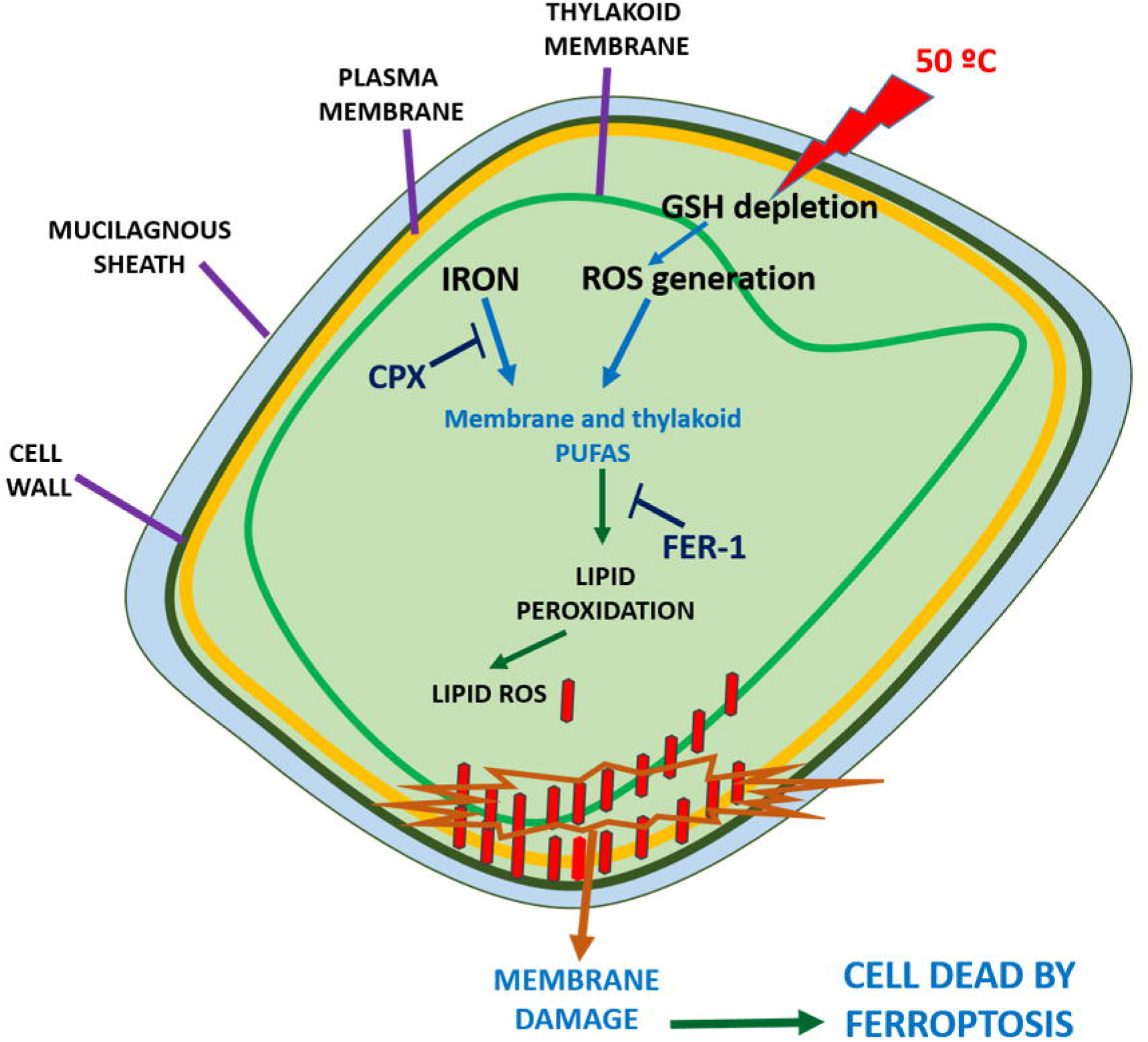
Schematic view of the ferroptotic cell death pathway triggered by heat shock in *Synechocystis* sp. PCC 6803 cells. Early GSH depletion following the heat shock, leads to lipid peroxidation, mostly of polyunsaturated fatty acids from thylakoid and plasma membranes. CPX or Fer-1, canonical ferroptosis inhibitors,, prevent the accumulation of toxic ROS, inhibiting ferroptotic cell death.

Altogether, these findings open up a broad field of future research investigating cell death in cyanobacteria, which could be of great relevance in the management of toxic blooms and their ecological consequences.

## Materials and methods

### Cyanobacterial cells and culture conditions

Axenic cultures of *Synechocystis* sp. PCC 6803 were grown in an orbital shaker (120 rpm) at 28 ± 2°C, under constant light (30 μE m^−2^ s^−1^) in BG11 medium buffered with 20 mM HEPES-KOH to pH 7.5 (Rippka et al., 1979).

### Pre-incubations and heat treatments

Cultures growing in control condition were submitted to heat treatments in a water bath when they reached logarithmic phase (approximately OD_750_ 0.8-0.9). Cells was exposure to50°C or 77°C for different times (10, 30, 60 min, 2 h, 4 h, 6h). GSH (L-Glutathione reduced form, Sigma; final concentration 100 μM), AsA (L-ascorbic acid, Merk; final concentration 1 μM), D-PUFAs (Retrotope; final concentration 50 μM), CPX (final concentration 1 μM) and Fer-1 (final concentration 1 μM) were applied to *Synechocystis* sp. PCC 6806 cultures 24 h before the HS. A second pulse of Fer-1 (1 μM) was added 2 h before the HS. For Cl_2_Ca and EGTA experiments, compounds were applied as follows: a) Cl_2_Ca (3 mM final concentration) 24 h before HS, b) EGTA (1mM final concentration) 24 h before HS; c) Cl_2_Ca (3 mM) for 2 h followed by addition of EGTA (1mM) 22 h before HS (Distéfano et al., 2017). d) EGTA for 2 h followed by incubation with fer-1 or CPX for 22 h before inducing cell death by treating cells at 50°C for 4 h. Cultures with no addition were used as controls.

### Drop tests for viability analysis

The effect of H_2_O_2_ (2, 5 and 10 mM, for 1 h) and HS (50°C and 77°C, for 10 min) in the absence or presence of Fer-1, CPX, GSH, AsA, EGTA, CaCl_2_ was tested on solid medium. Serial dilutions of treated cultures were prepared (10^0^ to 10^−3^) and spotted onto solid BG11 agar plates and cultivated under constant light (30 μE m^−2^ s^−1^) at 26 ± 2°C, for 20 days.

### Cell counting and viability determination

*Synechocystis* cell suspensions were stained with SYTOX Green at final concentration of 1 μM for 10 min protected from light. Cells were examined and counted by light and fluorescence microscopy using a Nikon E600 microscope equipped with a B-2A cube with 450–490 nm excitation and 500–515 nm emission filters, and a Neubauer chamber. Images were captured by Olympus DP72 digital camera, using Cellsens Entry imaging software. At least 10 random fields were taken for viability calculations in each experiment and quantified in Image J (https://imagej.nih.gov/ij/) (Schulze et al., 2011) (Figure S2 a and b).

### TEM analysis

*Synechocystis* cells subjected to H_2_O_2_ (10 mM for 1 h) and HS (50°C and 77°C for 10 min) were fixed with 2.5% (vol/vol) glutaraldehyde in PBS buffer overnight at 4°C. After washing three times in PBS buffer, samples were post fixed with 1% (vol/vol) osmium tetroxide in medium buffer for1 h and washed twice in distilled water. Samples were dehydrated in increasing concentrations of alcohol and embedded in Spurr epoxy. Ultrathin sections (90 nm) were stained in uranyl acetate and lead citrate and examined with a JEM 1200 EX II transmission electron microscope (JEOL Ltd). Images were captured using a digital camera (Erlangshen ES 1000 W, model 785) from the Central Service of Electron Microscopy of the Faculty of Veterinary Sciences, Universidad Nacional de La Plata (Argentina).

### Measurement of glutathione and ascorbic acid content

Fifty milliliters of exponentially-growing culture (∼2.5 × 10^7^cells ml^−1^) were collected by centrifugation (9,300 × g for 10 min, 4 °C), re-suspended in 1 ml 3% trifluoracetic acid, mixed thoroughly (vortex for 1 min), freezed with liquid nitrogen, thawed in ice and mixed in vortex for 1 min prior centrifugation (16,000 × g 15 min, 4 °C). Supernatants were passed through a C-18 column (Bond Elute, Varian) for a partial sample purification and eluted in 1 ml 100 mM phosphate buffer pH 7. Then supernatant was used for GSH and AsA determinations.

The determination of reduced AsA and dehydroascorbate (DHA) (the oxidized form of AsA) was carried out following Bartoli et al. (2006). The content of AsA was determined with a HPLC system using an UV-VIS detector (Model SPD-10AV, Shimadzu®, Japan) at λ = 265?nm coupled with a LC-10 AT pump (Shimadzu®, Japan) and compounds separated in a C-18 column (Microsphere C-18 SS 100 × 4.6?mm, Varian Inc., USA). Results were expressed in pmol of ascorbic acid per cell.

Determination of glutathione (GSH) and glutathione disulfide (GSSG) was done following the DTNB method using glutathione reductase and 2-vinylpyridine as described by Griffith (1980). Results were expressed in μmol of GSH or GSSG per cell per mg of fresh weight basis, and in nmol per cell.

### Flow cytometry experiments

Cell viability, ROS production, lipid peroxidation and caspase-like activity were assessed by flow cytometry in a Partec Cyflow Space cytometer equipped with a 488 nm laser (blue) and 3 detectors: 525 nm (15 BP) (green), 590 nm (25 BP) (orange) 675 nm (10 BP) (red). *Synechocystis* cells in logarithmic phase (OD = 0.8) were used in all the assays.

To test viability, fluorescein diacetate (FDA) was used as it enters and emits fluorescence only in living cells (Gumbo et al., 2014). A FDA stock solution (1000X) was prepared by solubilizing 50 mg of FDA in 5 ml of DMSO and stored in the dark at − 20 C until further use. Cultures pre-treated or not with Fer-1 and CPX for 24 h, were subjected to 50°C and 77°C for 10 min. After 16 h, cultures were incubated with FDA for 10 min. General cytosolic ROS was measured using H_2_DCFDA. A 10 mM H2DCFDA stock solution was prepared in DMSO and stored in the dark at −20°C BODIPY® C-11 581/591 (undecanoic acid) was used for the analysis of lipid peroxidation. This probe detects the oxidation of the polyunsaturated butadienyl portion, changing its emission peak from ∼ 590 nm to ∼ 510 nm (Cheloni and Slaveykova, 2013). A 2mM (1000X) BODIPY stock was prepared and frozen at −20°C. Cultures pre-treated or not with Fer-1 or CPX for 24 h were subjected to 50°C and 77°C for 10 min. For ROS and lipid peroxidation analysis, aliquots were taken at different times (30 min, 1 h, 2 h, 3 h) after HS, incubated with H_2_DCFDA or BODIPY for 30 min and analyzed in the cytometer. Aliquots treated with 500 mM H_2_O_2_ were used as a positive control for the formation of ROS and oxidized lipids. Caspase-like activity was measure with CellEvent™ Caspase-3/7 Green Flow Cytometry Assay Kit (Invitrogen), a nucleic acid-binding dye that harbors the caspase-3/7 cleavage sequence, DEVD, and is fluorescent after being cleaved and bound to DNA. Analysis of raw flow cytometry data was done with FlowJo (https://www.flowjo.com/).

### RNA isolation and quantitative real-time RT-PCR

Target genes were identified in the genome of *Synechocystis* (http://genome.annotation.jp/cyanobase/*Synechocystis*) and PCR primers were designed with Primer-BLAST (www.ncbi.nlm.nih.gov/tools/primer-blast) (Table S1). The analyses of gene expression was performed by real-time quantitative PCR (RT-qPCR). Total RNA was extracted from cell pellets using TRIzol Reagent (Invitrogen) according to the manufacturer’s instructions. After digestion with RQ1 RNase-free Dnase (Promega), RNA (1 μ g) was retro-transcribed using random hexamers (Promega). The qRT-PCR reactions were performed in a Step One real time PCR system (Applied Biosystems) using a Micro Amp Fast Optical 48-well reaction plate with 15 μ L reaction volume containing 1 x Power Sybr Green PCR Master Mix (Thermo Fisher), 0.2 μM of each primer and 1.5 μg of cDNA. Cycling program was: one cycle of 95°C for 10 min, 40 cycles of 95°C for 15 s followed by 40 cycles of 54°C for 1 min. A melting curve analysis was conducted to verify the formation of a single unique product and the absence of potential primer dimerization. The different biological samples were subsequently normalized against expression of *rnpB* gene coding for the RNA subunit of RNaseP.

## Data analysis

All experiments were performed at least in triplicate. The results are expressed as the mean ± standard deviation (SD). The main effects of treatments were examined by running two-way ANOVA for AsA and GSH data, and two-way ANOVA in Generalized Linear Models (GLM) module for the rest of experiments. Binomial distribution was fitted using the *glm* function in R. When significant differences (p < 0.05) were found, Tukey post hoc test was used for multiple comparisons within groups. Statistical analyses were performed using the open access software R (R Core Team, 2018). The ANOVAs of the GLM models and post hoc comparisons were performed with *car* and *lsmeans* packages (Fox and Weisberg, 2010; Lenth, 2016)

## Online supplemental material

Suplementary figure 1. **(a)** Representative fluorescence images showing cell death in *Synechocystis* sp. PCC 6803 detected by SYTOX® Green nucleic acid stain. (b) Neither Fer-1 nor CPX prevented cell death triggered by H_2_O_2_ treatments. (c) Kinetics of cell death induced by 50°C. (d) GSH (100 μM) and AsA (1 μM) addition prevents cell death induced by 50°C HS (10 min). (a-d) Cultures were preincubated with DMSO, Fer-1 (1 μM), or CPX (1 μM) for 24 h before HS. (b and d) Viability was tested via drop test.

Suplementary figure 2: Caspase-like activity measured by flow cytometry using CellEvent after 50°C, 77°C for 10 min or with H_2_O_2_ 10 mM for 1 h.

## Acknowledgments

We thank Mikhail S. Shchepinov for providing D-PUFAs, Silvana Colman, Macarena Perez-Cenci and Viviana Daniel for technical assistance, Daniela Sueldo, and Juan José Guiamet for insightful comments.

This research was funded by grants to M.V. Martin from Agencia Nacional de Promoción Científica y Técnica Argentina (PICT 1956 and PICT 0173) and to

G.C. Pagnussat from Agencia Nacional de Promoción Científica y Técnica Argentina (PICT-2017-0201). A. Aguilera is a postdoctoral fellow of Consejo Nacional de Investigaciones Científicas y Técnicas (CONICET); F. Berdun is a fellow of Consejo Interuniversitario Nacional. M.V. Martin, G.C. Pagnussat, G. Salerno G. and C.G. Bartoli, are CONICET researchers.

The authors declare no competing financial interests.

## Author contributions: Author contributions

Conceptualization: M.V. Martin, G.C. Pagnussat, A. Aguilera, F. Berdun, G. Salerno; Methodology: M.V. Martin, G.C. Pagnussat, A. Aguilera, F. Berdun; C. Bartoli; Charlotte Steelheart, Alegre, Matías Leonel; Investigation: M.V. Martin, A. Aguilera, F. Berdun, G.C. Pagnussat, G. Salerno Formal analysis: M.V. Martin, A. Aguilera, F. Berdun, G.C. Pagnussat; Supervision: M.V. Martin, G.C. Pagnussat; Project administration: M.V. Martin, G.C. Pagnussat; Validation: M.V. Martin, G.C. Pagnussat, A. Aguilera, F. Berdun; Resouces: M.V. Martin, G.C. Pagnussat, G. Salerno; C. Bartoli; Funding acquisition: M.V. Martin, G.C. Pagnussat; Visualization: M.V. Martin, G.C. Pagnussat, A. Aguilera; Writing of the original draft: M.V. Martin, G.C. Pagnussat, A. Aguilera, F. Berdun; G. Salerno; Writing, review and editing: M.V. Martin, G.C. Pagnussat, A. Aguilera, F. Berdun, G. Salerno; C. Bartoli, Charlotte Steelheart.

