## Supplementary figures and images for "Heat stress induces ferroptosis in a photosynthetic prokaryote"

### Supplemental figure 1

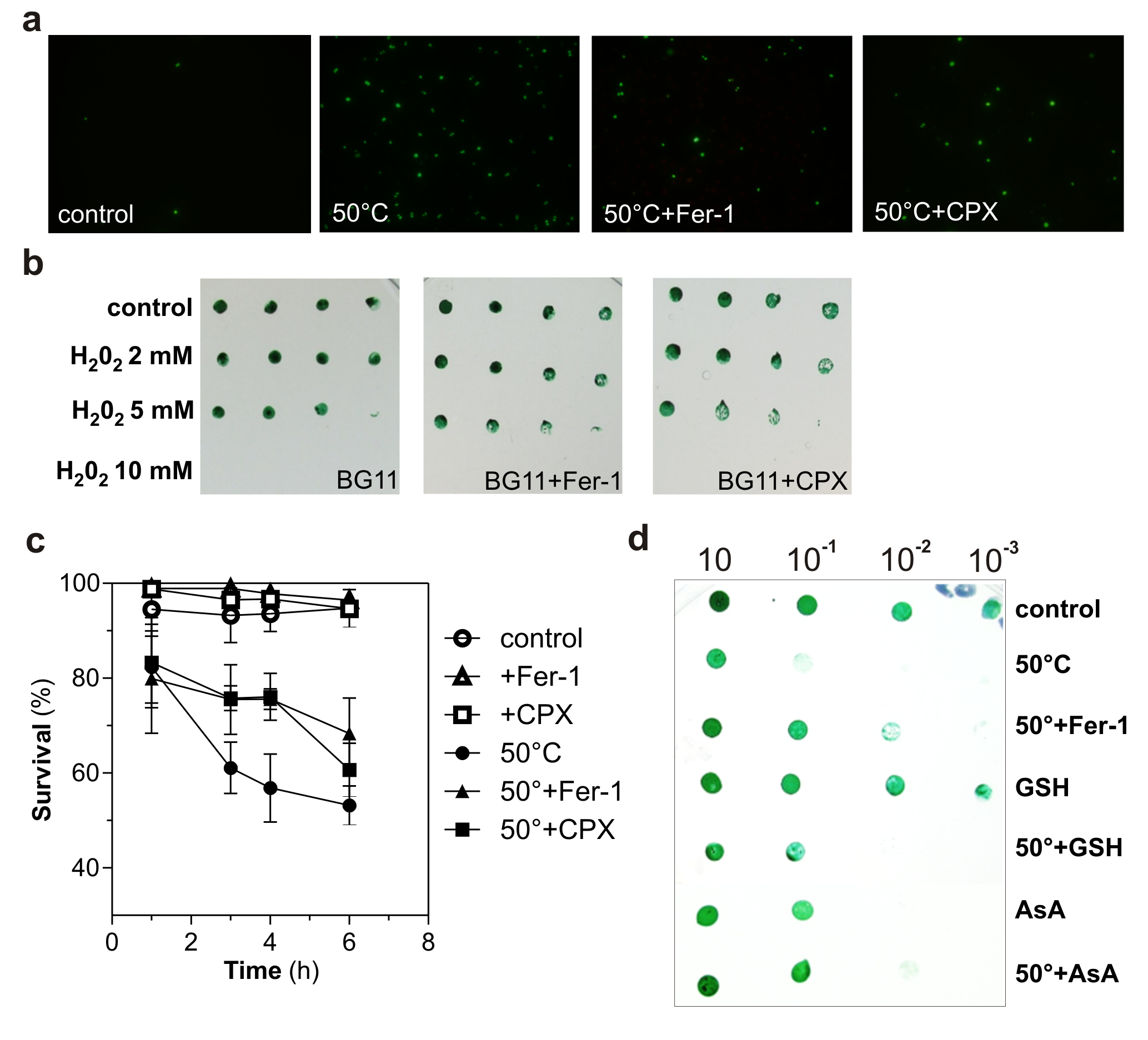

### Supplemental figure 2

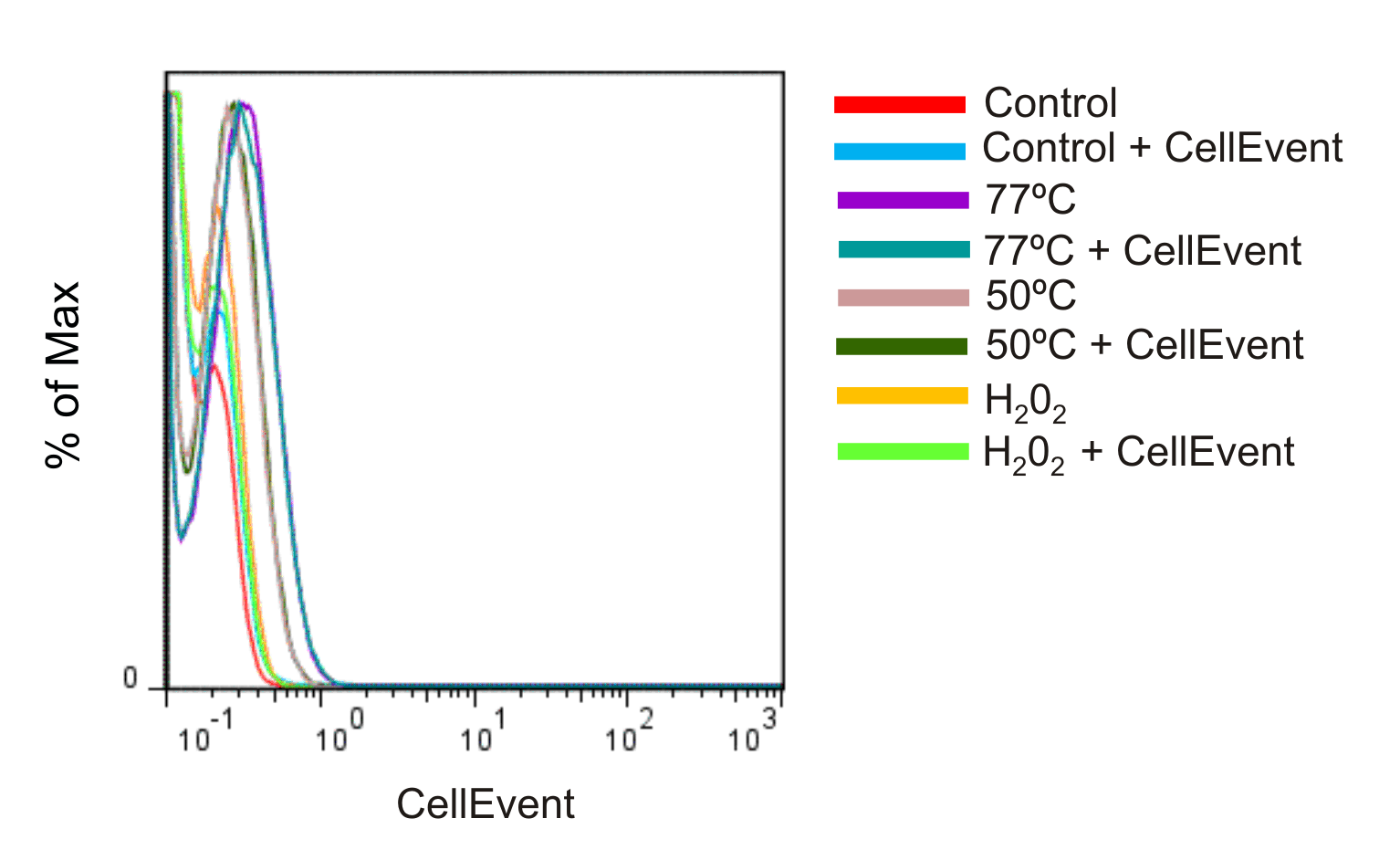
