## Supplemental Table 1 for "Heat stress induces ferroptosis in a photosynthetic prokaryote"

**Supplementary table 2:** Primers used in this study in qPCR reactions. Primers were designed using sequences corresponding to genes identified in the genome of *Synechocystis* sp. PCC 6803 (http://genome.annotation.jp/cyanobase/Synechocystis)

| Gene | Locus tag | Sequencing primer 5´-3´ | Product size (bp) |
|---|---|---|---|
| *gpx1* | slr1171 | GCCAACAACACAATCTACGG<br>GAGTGAAGCCACATTGACTG | 120 |
| *gpx2* | slr1992 | CCCTCTACCAATTCCTTGTG<br>CCTTGAGATTAGTGTCATCGG | 131 |
| *gshA* | slr0990 | CCTTACCATACCTACATCGAACAA<br>GGCTTCCATCCTCACTAAAC | 125 |
| *gshB* | slr1238 | GAAGGAAGGTGATAAGCGAAT<br>CTGGAGGTAATGGTAGTGGG | 134 |
| *ggt* | slr1269 | AAAGTATTGATTGAACCAGCGG<br>AAACTTTCTGCCTTCCTCTTG | 122 |
| *groEl1* | slr2076 | GTATTGATAAGGCCACTGATTTCC<br>CCAACTTCTTCGTCGTTACC | 120 |
| *groES* | slr2075 | TTGCCTGATAACGCCAAAGA<br>GAATAGAGAACCTTATCGCCCAC | 124 |
| *furA* | sll0567 | CGATTCCCTCAAAGCCGAAC<br>TCTTCTTCCAGACGATGGTGTA | 137 |
| *feoB* | slr1392 | GCGATCACCCTCTTAAACCT<br>GACTACAACCCGTTCCGTAT | 129 |
| *futA* | slr1295 | TATATTCATCCCGGCACTACAA<br>GTTCGTCTGCCTTACCTTCAA | 111 |
| *futB* | slr0327 | TGTCCTATCCCTACGTTTACCT<br>TAAAGCCACCCGACTGAAAC | 121 |
| *futC* | sll1878 | CCGCCTTTCCATAGAAGACT<br>CGATGGTCCTAATAGTCCCAG | 135 |
| *rnpB RNase P subunit B* | slr1469 | GTGAGGACAGTGCCACAGAA<br>CCTTTGCACCCTTACCCTTT | 120 |
